# Arrhythmia Mechanisms and Spontaneous Calcium Release: II - From Calcium Spark to Re-entry and Back

**DOI:** 10.1101/434456

**Authors:** Michael A. Colman

## Abstract

**Motivation:** The role of sub-cellular spontaneous calcium release events (SCRE) in the development of arrhythmia associated with atrial and ventricular tachycardia and fibrillation has yet to be investigated in detail. SCRE may underlie the emergence of spontaneous excitation in single cells, resulting in arrhythmic triggers in tissue. Furthermore, they can promote the substrate for conduction abnormalities. However, the potential interactions with re-entrant excitation have yet to be explored. The primary aim of this study was therefore to apply a novel computational approach to understand the multi-scale coupling between re-entrant excitation and SCRE.

**Methods:** A general implementation of Spontaneous Release Functions - which reproduce the calcium dependent SCRE dynamics of detailed cell models at a significantly reduced computational cost - was used to reproduce SCRE in tissue models. Arrhythmic dynamics, such as rapid pacing and re-entry, were induced in the tissue models and the resulting interactions with SCRE were analysed.

**Results:** In homogeneous tissue, the emergence of a spontaneous beat from a single source was observed and the positive role of coupling was demonstrated. Conduction block could be promoted by SCRE by both inactivation of the fast sodium channel as well as focal pacing heterogeneity interactions. Sustained re-entrant excitation promoted calcium overload, and led to the emergence of focal excitations both after termination of re-entry and also during re-entrant excitation. These results demonstrated a purely functional mechanism of re-entry and focal activity localisation, related to the unexcited spiral wave core.

**Conclusions:** SCRE may interact with tissue excitation to promote and perpetuate arrhythmia through multiple mechanisms, including functional localisation and mechanism switching. These insights may be particularly relevant for successful pharmacological management of arrhythmia.

## Introduction

Tachycardia and fibrillation describe rapid cardiac arrhythmias which interrupt the regular electrical activity of the heart and can lead to reduced cardiac output and sudden cardiac death [1]. The underlying rapid and irregular electrical activation of cardiac tissue may be mediated by abnormal spontaneous pacing (focal ectopic activity), self-perpetuating re-entrant excitation, or a complex interplay between both mechanisms [1,2]. Management of these arrhythmias is typically challenging, often requiring invasive procedures such as implanted defibrillators or catheter ablation; even these interventions have limited success rates [3,4]. Improved understanding of the mechanisms underlying the genesis, perpetuation and recurrence of rapid arrhythmias is vital for the development of improved treatment strategies.

The potential links between stochastic sub-cellular dynamics of the intracellular calcium (Ca^2+^) handling system and pro-arrhythmic cellular electrical phenomena have been highlighted by multiple research groups [5-9], yet the contributions and mechanisms of their manifestation at the organ scale as arrhythmia have yet to be described in detail. This manuscript accompanies a preceding study, Arrhythmia Mechanisms and Spontaneous Calcium Release: I - Multi-scale Modelling Approaches (Under Review), which outlines in detail the multi-scale mechanisms of spontaneous Ca^2+^ release events (SCRE) and an approach to overcome the challenges of computational modelling these phenomena at the organ-scale; the reader is therefore referred to that manuscript for detail, and only a brief summary of the relevant mechanisms follows.

A feedback mechanism emerging from the structure-function relationships underlying cardiac cellular excitation-contraction coupling presents a potential pro-arrhythmic pathway for the propagation of random state-transitions at the microscopic scale (sub-cellular) to the macroscopic (whole-organ): (i) spontaneous opening of the ryanodine receptors (RyRs), which control release of Ca^2+^ from the Sarcoplasmic Reticulum (SR) into the intracellular space, can trigger further openings within the dyad (spatially restricted sub-spaces to localise the SR and sarcolemmal membranes) and lead to a Ca^2+^ spark; (ii) the spatially distributed dyads present a sub-cellular substrate for the propagation of Ca^2+^ sparks as a whole-cell Ca^2+^ wave; (iii) this spontaneous whole-cell Ca^2+^ transient can lead to cellular delayed-after-depolarisations (DADs) and triggered action potentials (TA) through activation of the sodium-calcium exchanger, NCX; (iv) TA may propagate in tissue as ectopic focal excitation, potentially leading to the initiation of transient or sustained arrhythmia.

Recently, a few computational studies have demonstrated the mechanisms by which SCRE may emerge as focal excitation and conduction block [10–16]. However, there is still much to be understand about the role of SCRE in cardiac arrhythmia, for example related to its long-term interactions with re-entrant excitation.

In this manuscript, the multi-scale framework presented in the preceding study is applied to: (i) highlight the mechanisms of synchronisation of SCRE to form an ectopic beat, and (ii) investigate the role of SCRE variability in its SR-Ca^2+^ dependence; (iii) demonstrate different mechanisms of SCRE mediated conduction block; and (iv) investigate the coupling between SCRE and re-entry underlying sustained arrhythmia. The results illustrate the general mechanisms of coupling between SCRE and arrhythmia, present a novel mechanism of functional localisation of focal and re-entrant excitation, and represent an initial step towards full assessment of the importance and mechanisms of SCRE in cardiovascular disease morbidity and mortality.

## Methods

### Multi-scale modelling of spontaneous calcium release events

This section only briefly outlines the detailed methods presented in the preceding study, Arrhythmia Mechanisms and Spontaneous Calcium Release: I - Multi-scale Modelling Approaches (Under Review); Full model equations are also provided in the Supplementary Material S1 Text (Model Description).

#### Computational Framework

The basis of the computational framework is a structurally idealised model of spatio-temporal calcium handling which is capable of reproducing SR-Ca^2+^ dependent SCRE (Figure 1A). The distributions of the initiation time, duration and peak of open RyRs waveforms - the parameter which controls SCRE morphology - can be described by a small number of parameters (Figure 1B); These parameters can thus be defined as functions of SR-Ca^2+^ (Figure 1C) and individual waveform parameters - which describe the shape of analytical Spontaneous Release Functions (SRF; Figure 1D) - can be randomly sampled from the appropriate inverse functions in order to generate SCRE which vary in timing and morphology with the given SR-Ca^2+^ dependence. These SRF can then be incorporated into Hodgkin-Huxley type non-spatial cell models and in idealised and realistic tissue models, allowing simulations of SCRE at the tissue scale (Figure 1E).

**Figure 1:**
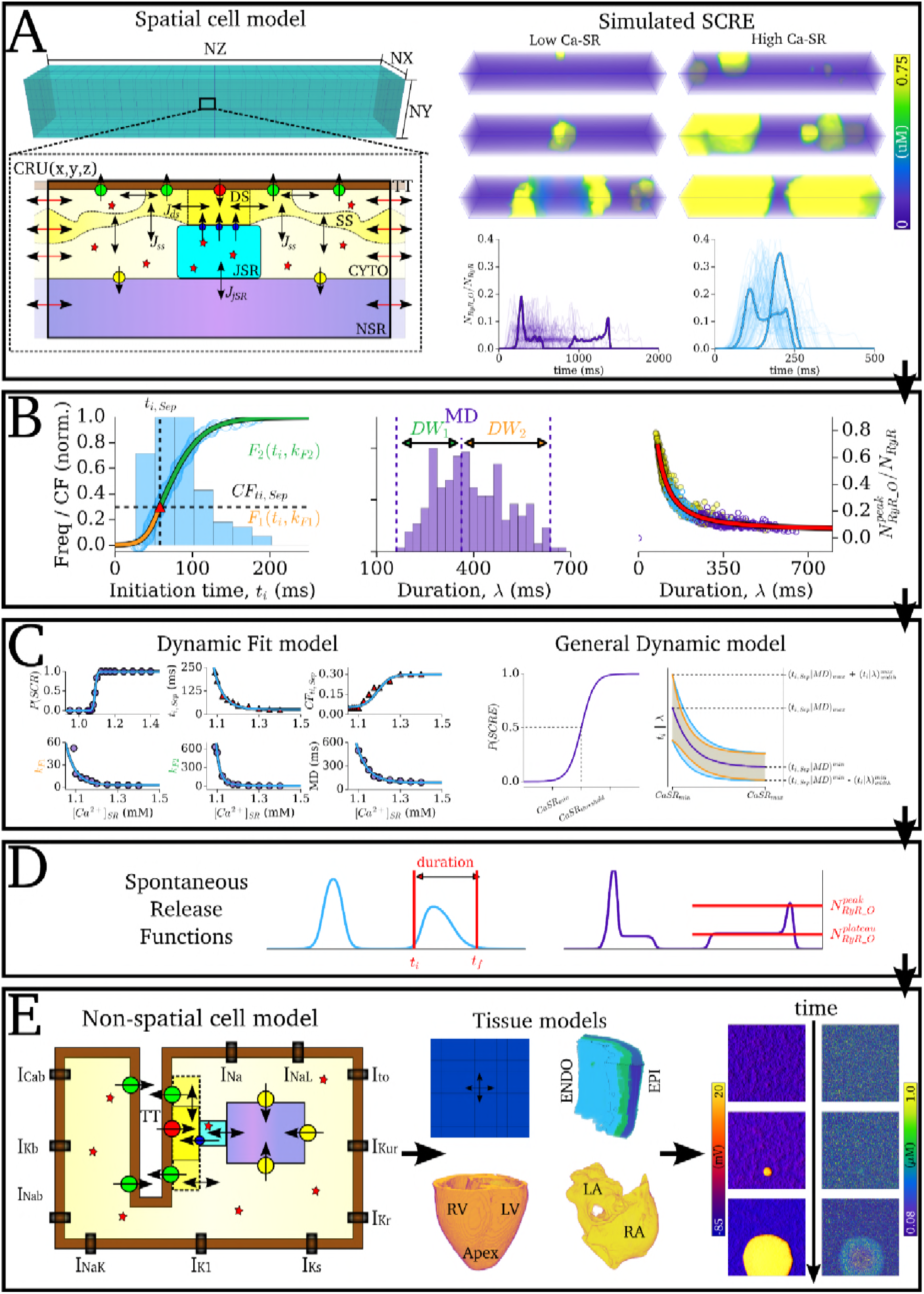
The computational framework for multi-scale simulation of spontaneous calcium release. A - 3D, microscopic Ca^2+^ handling model (left), illustrating the 3D grid of calcium release units (CRUs; upper panel) and the compartments and Ca^2+^ fluxes within a single CRU (lower panel). Labelled are the dyadic cleft space (DS), sub-space (SS), bulk cytosolic space (CYTO), network and junctional SR spaces (NSR, JSR), and a T-tubule (TT); simulated SCRE (right) showing three snapshots of Ca^2+^ waves at low (left) and high (right) SR-Ca^2+^; lower panels are overlays of 100 simulations at each SR-Ca^2+^. B - Statistics of SCRE, showing the distributions associated with initiation time (left), duration (middle) and peak RyR (right). Examples of two functions - and their relevant parameters - to fit the cumulative frequency of initiation time are shown. C - Defining the distribution parameters based on SR-Ca^2+^, illustrating the Dynamic Fit model (left) and General Dynamic model (right). D - Illustration of the analytical spontaneous release functions, defined by the parameters randomly sampled from the initiation time and duration distributions. Those illustrated correspond to the highlighted waveforms in A. E - Schematic of the non-spatial cell model (left), illustration of the tissue models (middle), and example simulation of a SCRE mediated focal excitation (right).

#### Spontaneous Release Functions (SRF)

Simple analytical waveforms corresponding to the open RyR waveforms during SCRE in the 3D cell model formed SRF for implementation in the non-spatial cell models. Briefly, the waveforms for the open RyR (*N*_RyR_O_) can be well approximated with the simple function:

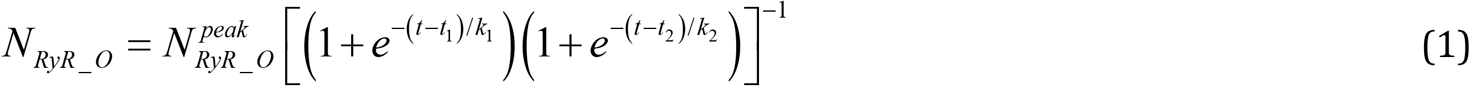

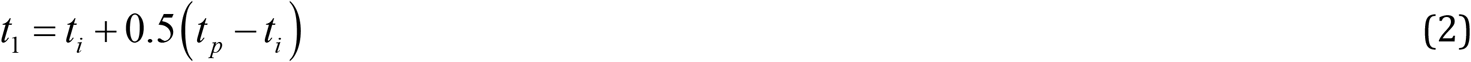

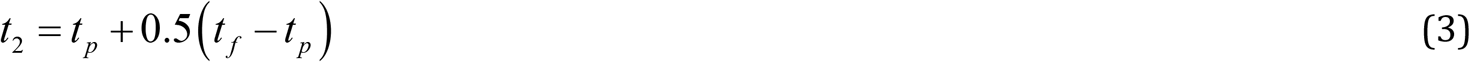

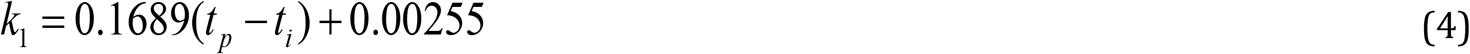

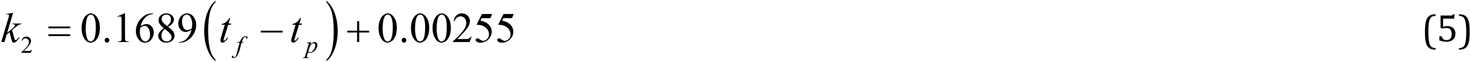

where *t_*i*_* is the initiation time of the SCRE, *t*_*f*_ is the end time (duration, *λ*, thus = *t*_f_−*t*_i_), *t*_p_ is the time of the peak of the waveform and *N*_RyR_O_^peak^ is the peak of open proportion RyR. The amplitude, *N*_RyR_O_^peak^, depends directly on the duration, and the peak time, *t*_p_, approximately varies evenly within the duration; the waveform is therefore completely described by two parameters: initiation time, *t*_i_ and duration *λ*. To reproduce stochastic variance, these parameters can be randomly sampled using the following inverse functions:

Initiation time, *t*_i_:

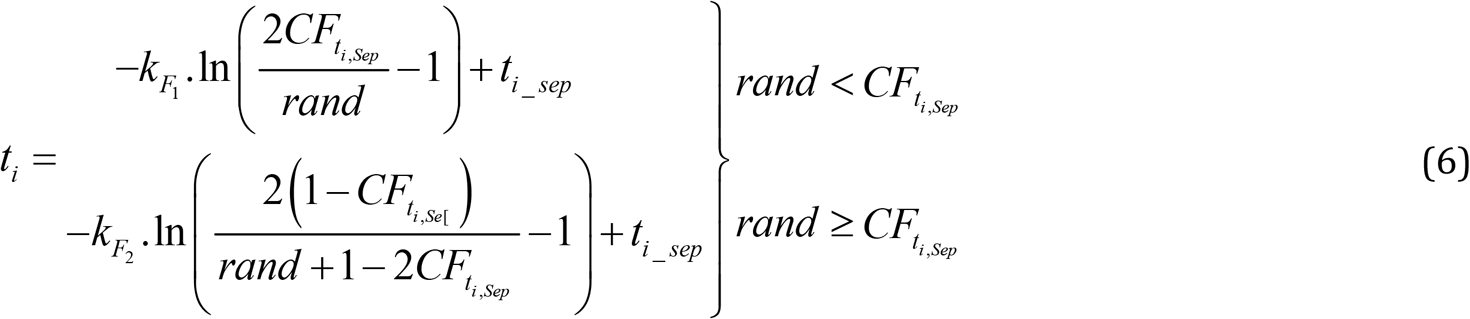

Duration, *λ*:

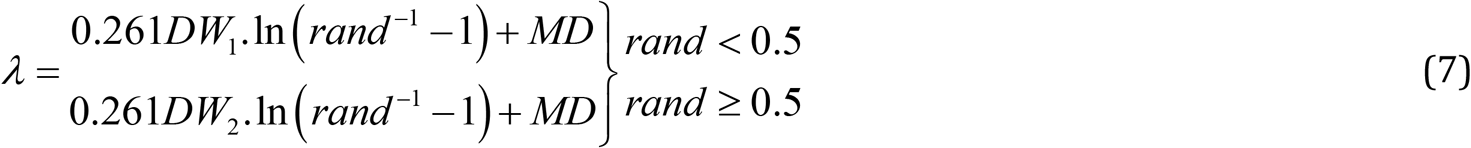

Where then peak time, *t*_p_, following a uniform distribution approximation:

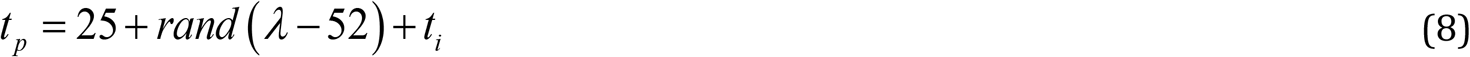

And the amplitude, *N*_RyR_O_^peak^:

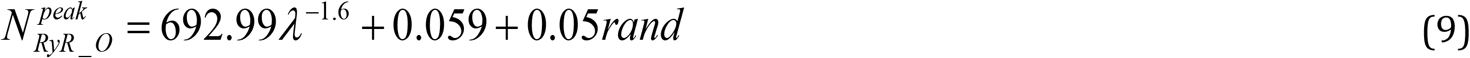

Where the parameters describing the initiation time and duration distributions (*t*_*i*,Sep_ - the initiation time corresponding to the point where two sigmoidal functions are separated; *CF*_ti,Sep_ - the cumulative frequency at this point (= *F*(*t*_i_)|_ti=ti,Sep_); *k*_F1_, *k*_F2_ - the gradient parameter of each function, corresponding to the width of the distribution either side of *t*_i,Sep_; *MD* - the median of the duration distribution; *DW*_1_, *DW*_2_ - the width of the duration distribution either side of the median, in ms) must be determined, in addition to the probability of release, *P(SCR)*.

Implementations were presented where these parameters were either directly set (corresponding to a specific dataset; Direct Control implementation), fit to the dynamics of the 3D cell model in multiple conditions (Dynamic Fit SRF implementation; Figure 1C), or where the SR-Ca^2+^-dependence of the distributions was determined through controllable inputs (General Dynamic SRF implementation; Figure 1C). Whereas a primary aim of the previous study was to develop an approach to allow direct translation of 3D cell modelling studies to the organ scale, the General Dynamic SRF implementation (Figure 1D) offers the most powerful approach to conduct the mechanistic investigation of this study’s ambitions. In this implementation, the SRF input parameters (*P*(SCR), *t*_i_sep_, *CF*_ti,Sep_, *k*_F1_, *k*_F1_, *MD*), which define the inverse functions from which the actual waveform parameters are sampled (equations 6-9), are determined from the SR-Ca^2+^ by the following functions:

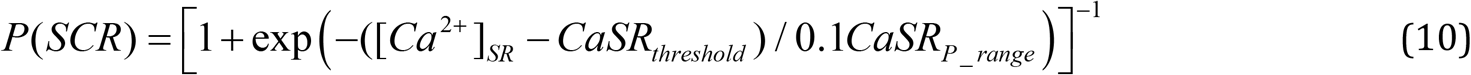

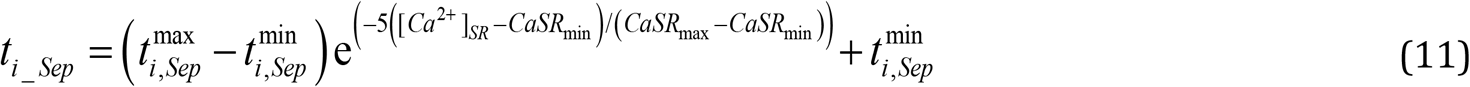

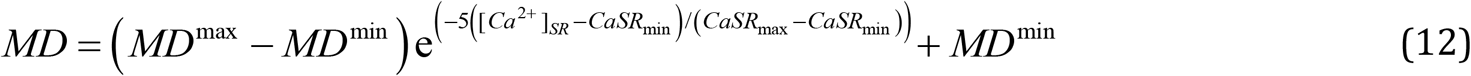

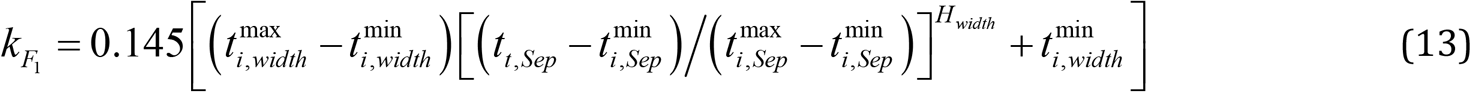

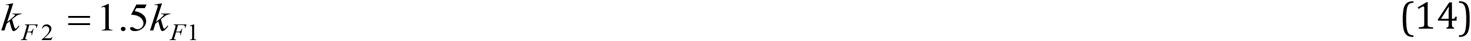

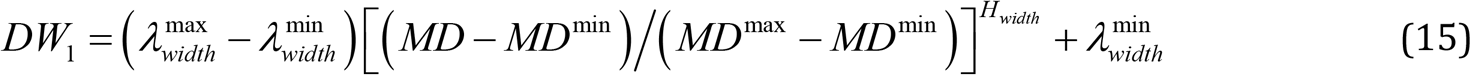

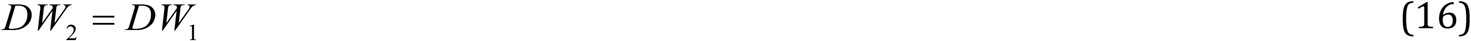

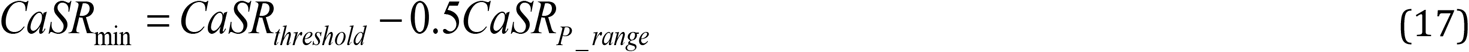

Where the primary variables describing the SR-Ca^2+^ dependence can be set as desired. The variables are: (i) - The threshold for SCRE (*CaSR*_threshold_); (ii) - The SR-Ca^2+^ range over which *P*(SCR) varies from 0 to 1 (*CaSR*_P_range_); (iii) - The maximal SR-Ca^2+^ above which SCRE distributions converge (*CaSR*_max_); (iv) - The minimum and maximum *t*_i,Sep_ and *MD* (*t* ^min^, *t* ^max^, *MD*^min^, *MD*^max^); (v) - The *t*_i_ and *λ* distribution widths at these extremes (*t*^min^, *t*^max^, *λ*^min^, *λ*^max^); And (vi) - the non-linearity of width variance (*H*_width_); *CF*_ti,Sep_ was set to 0.4.

### Simulation protocols

Focal excitation was initiated in tissue by pre-pacing the model at a rapid rate in combination with enhanced Ca^2+^-current and intracellular Ca^2+^ uptake activity, generally representing pro-SR-Ca^2+^ loading conditions observed with sympathetic stimulation (ISO) and disease remodelling (Supplementary Material S1 Text (Model Description)). The model was then left for a quiescent period, in which the impact of SCRE can be observed.

The General Dynamic SRF implementation was used to assess the relationship between SR-Ca^2+^ and focal excitation in tissue, allowing simulation of both homogeneous and heterogeneous media pertaining to the vulnerability to and dynamics of SCRE. A baseline SRF model was set with an SR-Ca^2+^ threshold for SCRE of 1.10 mM; four alternative models were introduced with this SR-Ca^2+^ threshold varied by ± 0.05 and 0.1 mM; all other parameters are either identical or set relative to the threshold (Supplementary Material S1 Text (Model Description)); Implementations with a lower SR-Ca^2+^ threshold exhibit larger amplitude and more rapidly timed SCRE than those with a higher SR-Ca^2+^ threshold at a specific concentration. A small amount of heterogeneity was first introduced by randomly assigning 60% of the cells to the baseline, and 10% to each of the four other distributions; a larger amount of heterogeneity was introduced by randomly assigning 20% of the cells to each of the five distributions.

Two potential mechanisms of SCRE-induced unidirectional excitation conduction block, which presents the possibility to degenerate into re-entry, have been shown in the studies of Liu et al., 2015 [15] and Campos et al., 2017 [11]: a well-established mechanism, in which focal excitation (in this instance as a result of SCRE) interacts with heterogeneity in refractory period and leads to asymmetric propagation patterns; and a novel mechanism by which DADs in combination with sodium channelopathy led to heterogeneous inactivation of the fast sodium current and conduction block. To demonstrate the capability of the SRF implementation to reproduce these dynamics, and to provide independent verification of the results in those previous studies, two protocols were used. DAD mediated conduction block was simulated in a homogeneous 2D sheet of the epicardial layer of the ventricle in combination with impaired sodium current (conductance of *I*_Na_ scaled by 30%; 5 mV shift of the inactivation steady-state); localised focal excitation was simulated in a 2D sheet model of the transmural celltypes in the ventricular wall (simple 1:1:1 ratio of EPI, M and ENDO cells), with the stimulus applied uniformly to the edge of the ENDO region. A small region was set to reduced *I*_K1_ to promote localized excitation. The Direct Control implementation was used to impose early-timed SCRE.

Re-entrant excitation was initiated using the phase-distribution method [17,18] applied to multiple cellular conditions, in general in combination with a reduction in the conduction velocity (***D*** reduced by up to half). In conditions in which re-entrant excitation sustained for more than 10-20 s, termination was induced by a decrease in the conductance of repolarising potassium currents and/or an increase in the conduction velocity (some conditions also spontaneously self-terminated after 10 s, primarily as a result of prolonged APD due to larger Ca^2+^ transients and therefore NCX activity).

## Results

In this section is described various applications of the multi-scale modelling framework, to: (i) demonstrate the mechanism of a stochastically mediated SCRE-induced ectopic beat; (ii) examine the complexities underlying the SR-Ca^2+^ dependence of focal excitation; (iii) demonstrate two independent mechanisms of conduction block associated with SCRE; and (iv) study the potential multi-scale, pro-arrhythmic interactions between SCRE and re-entrant excitation.

### Co-ordination of a stochastically initiated SCRE mediated focal excitation

Evaluation of the focal activation emerging in the 2D sheet illustrates the mechanism by which these small scale cellular events result in full tissue excitation and the dual role of electrotonic coupling (Figure 2; Supplementary Material S1 Video): SCRE occurring in independent cells initially causes only a small depolarisation of the cell membrane, repressed by electrotonic coupling (Figure 2C, purple trace); however, this depolarisation is spread to surrounding cells due the electrotonic coupling (Figure 2B); due to this depolarised resting potential, SCRE occurring later can act to further depolarise the surrounding tissue (Figure 2C, blue trace); once local tissue is sufficiently depolarised, the probability of DADs manifesting as TA significantly increases and appropriately-timed firing cells can much more easily initiate a focal excitation (Figure 2C, orange trace).

**Figure 2:**
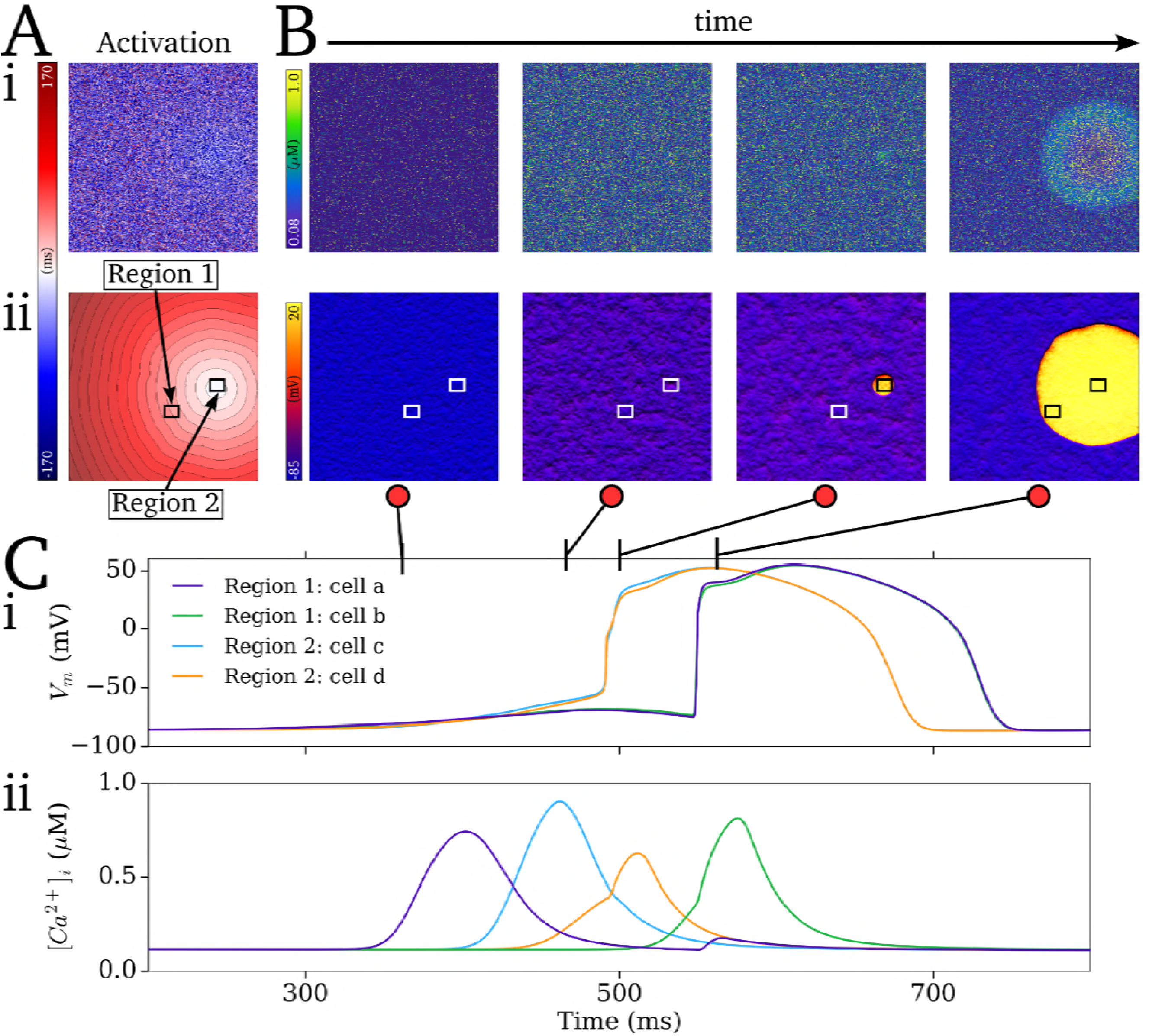
The role of electrotonic coupling in overcoming source-sink mismatch. A - Activation maps for intracellular Ca^2+^ (i) and membrane potential (ii) associated with a spontaneous focal excitation. Note that the time of the initiation of focal excitation (t = 0 ms) corresponds to the halfway point of the colour map. B - Temporal snapshots of Ca^2+^ (i) and *V*_m_ (ii) associated with the onset of focal excitation. C - *V*_m_ (i) and Ca^2+^ (ii) traces from individual cells from two regions with the tissue (labelled in Aii) illustrating the independent (Ca^2+^) and coupled (*V*_m_) cellular behaviour.

### Inter-cellular variability and the SR-Ca^2+^ dependence of ectopic activity

In homogeneous tissue, using a baseline General Dynamic SRF model parameter set (see Methods: Simulation Protocols), the probability curve for ectopic activity was substantially steeper than that for the emergence of TA from DADs in single cell (Figure 3Ai). Reduction of the density of *I*_K1_ shifted both the single cell and tissue TA probability curves to the left, towards the curve describing DADs (Figure 3Aii); however, the steep relationship in tissue remains.

With control *I*_K1_ density, both heterogeneity models (involving either 10% or 20% of cells randomly allocated each of the additional four General Dynamic SRF models with different SR-Ca^2+^ thresholds) shifted the SR-Ca^2+^ relationship to the right, despite the presence of cells more susceptible to SCRE; with reduced *I*_K1_ density, the small heterogeneity model still shifted the relationship to the right, but the large heterogeneity model now shifted it to the left (Figure 3B). In general, the introduction of heterogeneity reduced the steepness of the probability curve at low probabilities (Figure 3B). Heterogeneity models with the less vulnerable cells removed (higher SR-Ca^2+^ threshold cells reassigned to baseline) produced the same overall features: three cases still shifted to the right despite the presence of more vulnerable cells and absence of less vulnerable cells.

**Figure 3:**
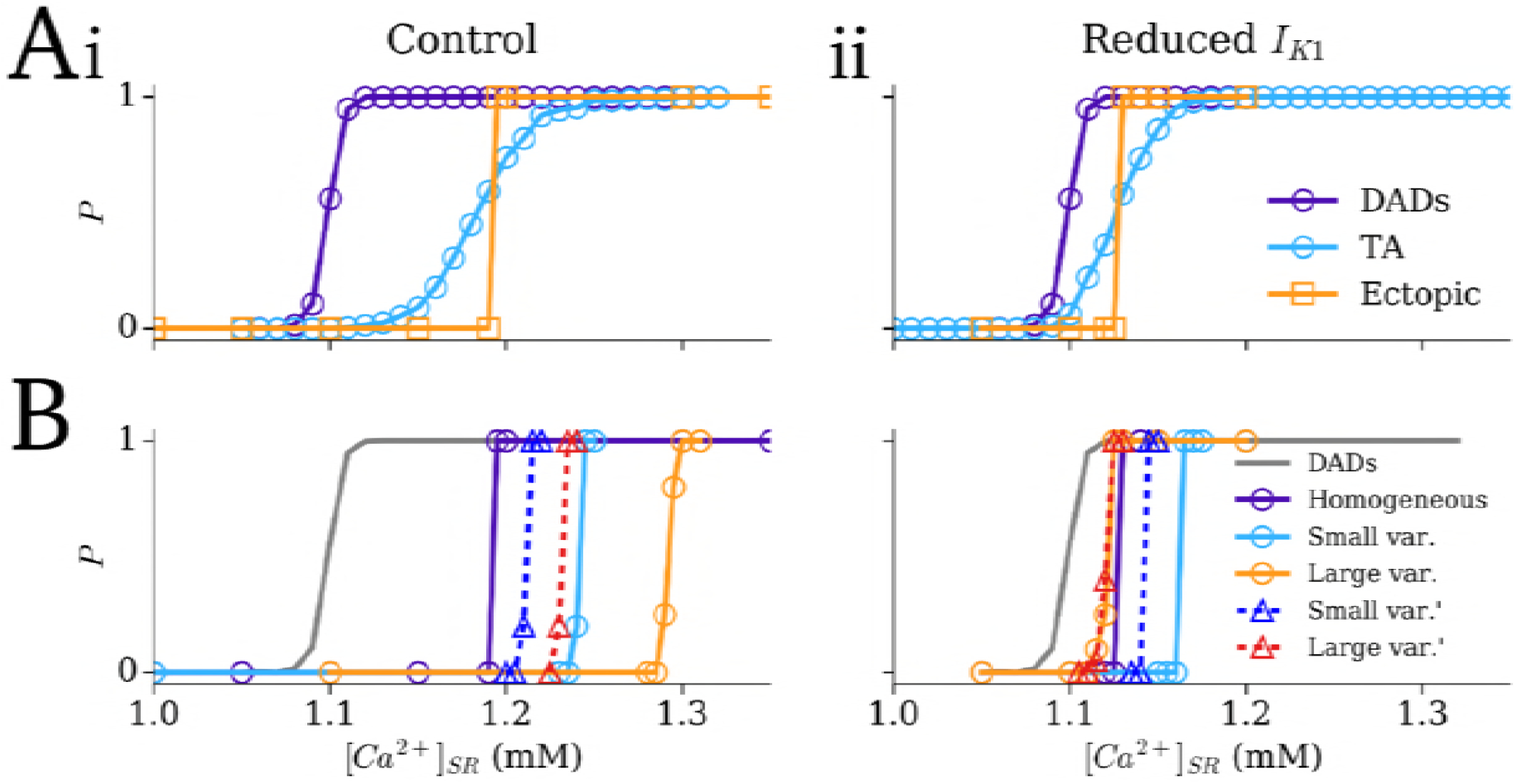
The effect of SCRE heterogeneity on the SR-Ca^2+^-TA relationship. A - Dependence of single cell DADs (purple), single cell TA (blue), and ectopic focal activity in 2D tissue (orange; square markers) on the SR-Ca^2+^ concentration in control (i) and reduced *I*_K1_ conditions (ii); homogeneous SCRE dynamics. B - Dependence of ectopic focal activity in tissue in the heterogeneous SRF conditions (purple; corresponding to the plots in A), with small and large variability (blue and orange), and with small and large variability with higher SR-Ca^2+^ threshold cells reassigned to baseline (blue and red; triangular markers); single cell DADs in the baseline condition are shown for reference (grey).

### Spontaneous Ca^2+^ release as a mechanism for conduction block

SCRE leading to non-TA inducing DADs was demonstrated to produce uni-directional conduction block during applied pacing in a homogeneous 2D model of the human ventricular wall (Figure 4A; Supplementary Material S2-S3 Video) under simulated sodium channelopathy conditions, wherein the SCRE induced DADs inactivate the sodium channel and result in non-uniform propagation following a stimulus uniformly applied to one edge (Figure 4Aii).

The potential for SCRE mediated focal excitation to result in uni-directional conduction block due to regional APD heterogeneity was investigated in the 2D transmural model of the human ventricular wall. Focal excitations originating in a small temporal window could exhibit a conduction block with the still refractory M-cell region; this was not observed in the homogeneous tissue (Figure 4B; Supplementary Material S4 Video). Note that the site of conduction block, unlike in simulations with an applied focal stimulus, is not necessarily clearly at the boundary between the EPI and M cells, but rather the focus emerges from the far side of the boundary.

**Figure 4:**
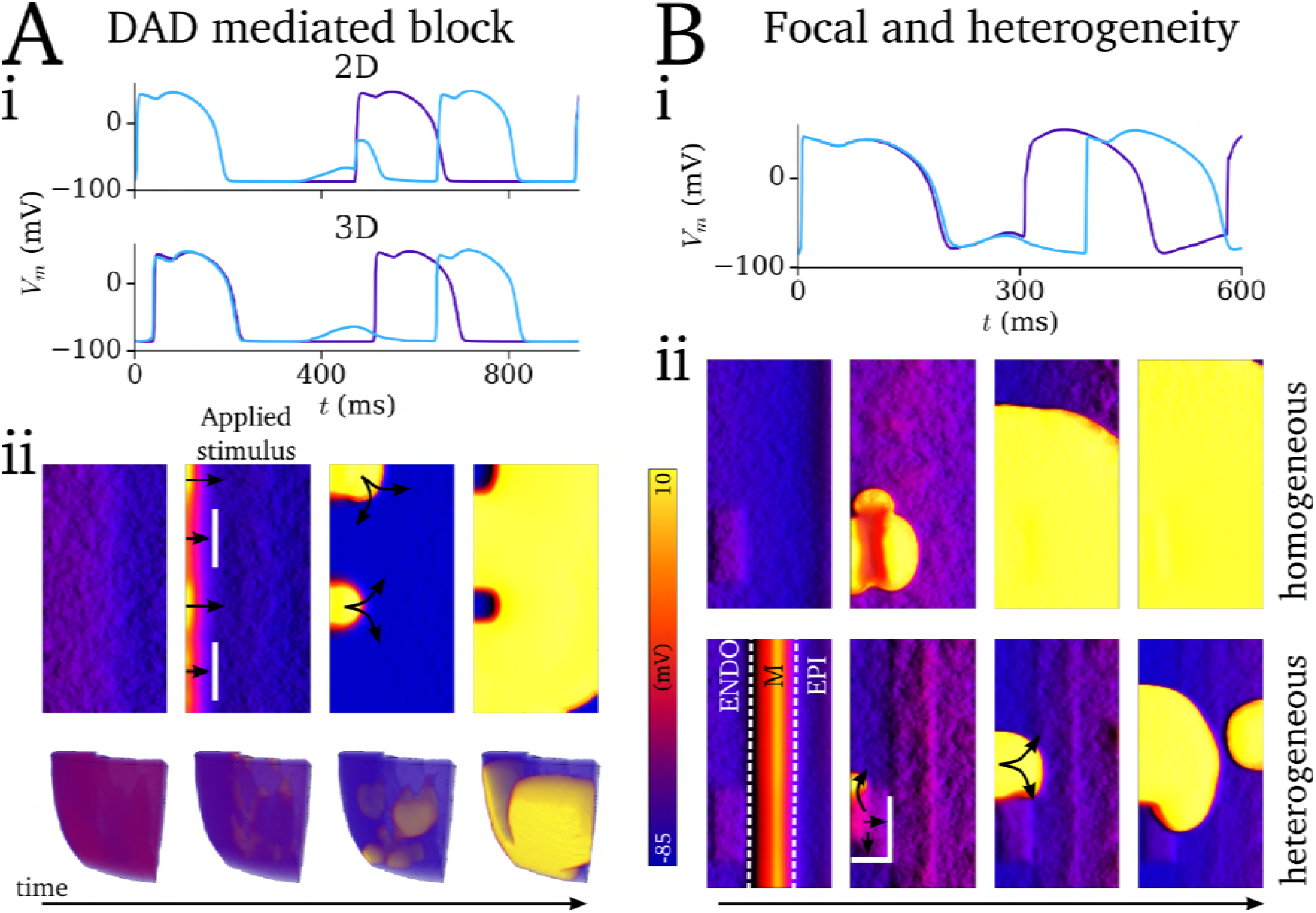
Mechanisms of SCRE mediated conduction block. A - AP traces (i) and temporal tissue snapshots (ii) of DAD mediated conduction block of an applied stimulus in 2D (upper panels) and 3D (lower panels) tissue simulations. In (i), the purple trace corresponds to simulations without SCRE and normal, successful propagation of the second AP (snapshots not shown) and the blue trace to the simulations illustrated in (ii), in which DADs inhibit the propagation of excitation following the applied stimulus. B - AP traces (i) and temporal snapshots (ii) of SCRE focal activity resulting in conduction block due to transmural AP heterogeneity in the ventricular wall. The purple trace in (i) corresponds to the homogeneous condition, in which the focal excitation propagates uniformly; the blue trace corresponds to the heterogeneous condition in which focal excitation propagates non-uniformly following conduction block.

### Re-entry promotes spontaneous Ca^2+^ release and focal excitation

Sustained re-entry combined with pro-arrhythmic conditions resulted in significant SR-Ca^2+^ loading (Figure 5Ai), which promoted the onset of SCRE mediated focal excitations following self-termination of re-entry (Figure 5Aii, B; Supplementary Material S5-S7 Video). Implementing a General Dynamic SRF model with the SR-Ca^2+^ threshold set below, but close to, the maximal SR-Ca^2+^ observed during re-entry (*CaSR*_threshold_ = 1.125 mM), a single, delayed focal excitation was observed following termination (Figure 5A, Bii); with a lower threshold (*CaSR*_threshold_ = 1.0 mM) multiple and rapid focal excitations were observed which perpetuated the arrhythmic dynamics for the duration of the simulation (Figure 5A, Biii). In general, where the tissue was sufficiently vulnerable for focal excitation to emerge, its origin was localised to the region of the scroll wave core on its final excitation pathway (Figure 5B; Supplementary Material S6-S7 Video); focal excitations emerging away from the scroll wave core were also observed closer to the threshold (Figure 5C).

**Figure 5:**
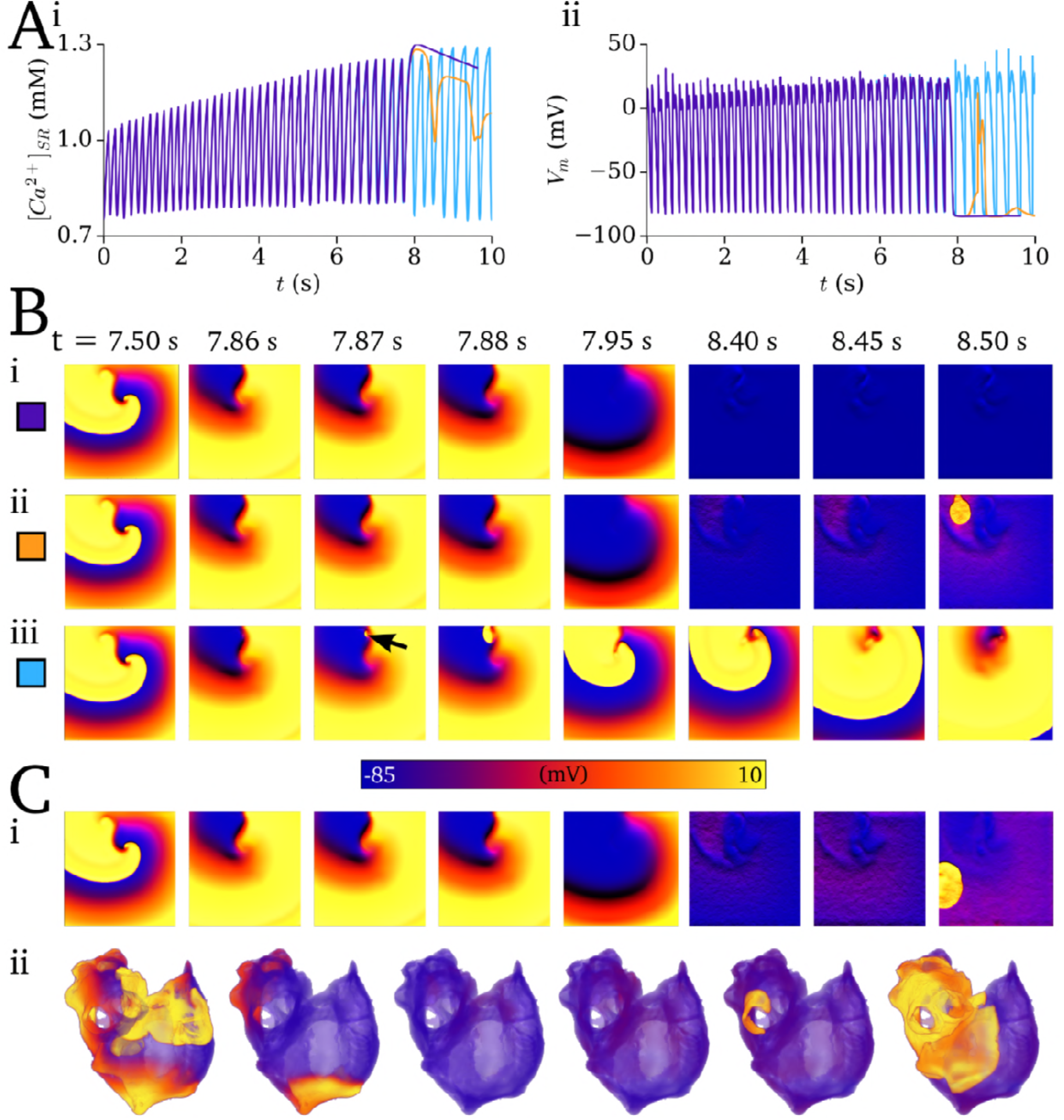
Coupling between re-entry and SCRE. A - SR-Ca^2+^ concentration (i) and *V*_m_ (ii) associated with sustained re-entry followed by self-termination (at around 8 s), simulations without SCRE (purple) and with the General Dynamic SRF model with two different thresholds (orange, 1.125 mM; blue, 1.0 mM). B - Temporal snapshots of voltage in the 2D sheet associated with the traces shown in A, showing self-termination (i) and the emergence of delayed (ii, corresponding to the orange traces in A) and rapid (iii, corresponding to the blue trace in A) focal excitations. C - Examples of non-localised focal excitations emerging in the 2D sheet (i) and 3D whole atria models (ii).

Localisation of focal excitation to the scroll wave core is determined by the dynamics of re-entrant excitation: the scroll wave core remains unexcited associated with each re-entrant cycle (Figure 6A), which leads to an island of high SR-Ca^2+^ in this region (Figure 6Ac). Due to this large SR-Ca^2+^ and longest recovery time, this region thus presents the earliest SCRE which may manifest as TA. This relationship is clearly illustrated by comparing the spatial distribution of recovery time (time since last AP) at the moment before focal excitation occurs with the associated focal activation map: the site of activation corresponds exactly (Figure 6Bi, iv-v) or approximately (Figure 6Bii- iii, vi) to the location of longest recovery time, in both single (Figure 6Bi-v) and double-scroll wave (Figure 6Bvi) simulations; the earlier the focal excitation occurs relative to the final re-entrant scroll wave, the stronger the correlation between longest recovery time and activation source and the more asymmetric the resulting activation pattern.

Under extreme pro-SCRE conditions, focal activity can occur at a rate comparable to re-entry; such pacing can transiently drive the excitation, potentially eventually terminating arrhythmia as the SR-Ca^2+^ depletes. Furthermore, the wavefront could degenerate back into a sustained re-entrant excitation (Figure 6C; Supplementary Material S8 Video). The latter is promoted by the asymmetric conduction patterns emerging from rapid focal excitation interacting with the tail of the previous scroll wave or asymmetric focal excitation. The underlying driving mechanism (focal or re-entrant) was not clear in AP traces from randomly selected cells in the 2D tissue, and could only be determined by spatio-temporally high resolution analysis of the excitation patterns (Figure 6C).

**Figure 6:**
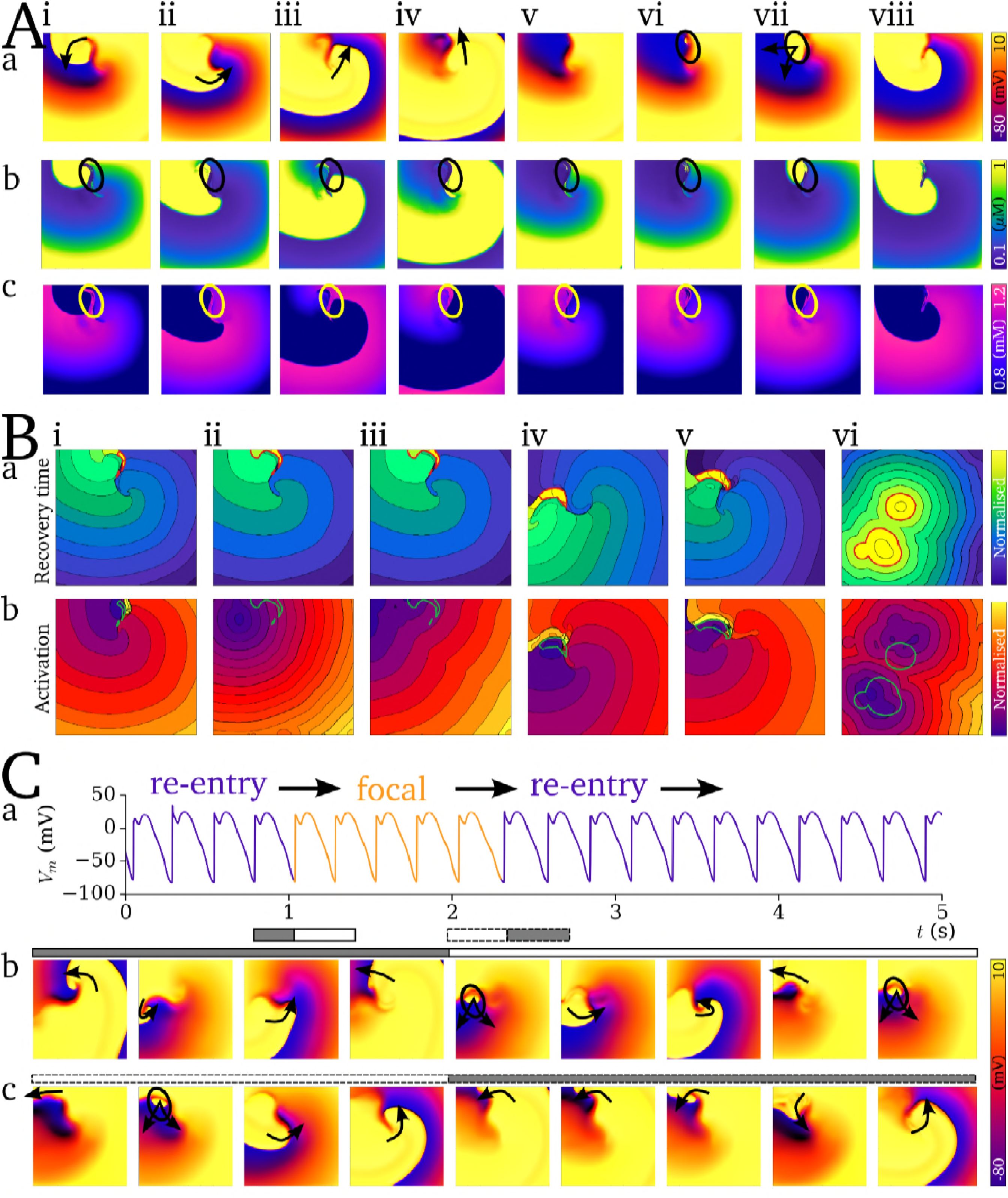
Functional localisation of re-entry and focal activity. A - Temporal snapshots (i-viii) of voltage (a), intracellular Ca^2+^ (b), and SR-Ca^2+^ (c) associated with the final, self-terminating re-entrant cycle and first focal excitation. Arrows in (a) indicate the conduction of the re-entrant and then focal excitations. Highlighted region (circle) illustrates the island of large SR-Ca^2+^ associated with the unexcited scroll wave core (i-iv) and its correlation with the focus of ectopic activation (vi-vii). B - Examples of recovery time maps (a) and focal activation maps (b) for 6 independent simulations (i-vi) associated with the self-termination of re-entry followed by ectopic excitation. The contour surrounding the region of longest recovery time (corresponding to the unexcited core illustrated in A) is highlighted in red in the recovery time map and green in the activation maps. Note the correlation between activation source and this highlighted region. C - Mechanism switching between re-entrant and focal excitation, showing the AP from a randomly selected cell (a) and temporal snapshots associated with the transition from re-entry to focal activity (b) and focal activity to re-entry (c). Snapshots corresponds to the temporal range illustrated by the grey and white bars with solid (re-entry to focal) and broken (focal to re-entry) borders.

## Discussion

### Summary

In this study, a novel computational framework for multi-scale modelling of SCRE was applied to study its involvement in arrhythmia mechanisms. Firstly, the dual role of electrotonic coupling in the emergence of SCRE as a full tissue focal excitation was illustrated (Figure 2): while initially repressing the magnitude of DADs, the small depolarisation can propagate through the tissue and allow later, appropriately localised and timed events to overcome the electrotonic load and activate an AP. Secondly, the role of cellular variability in SCRE dynamics on the SR-Ca^2+^ - focal excitation relationship was investigated (Figure 3), highlighting the complex constraints on timing, important role of *I*_K1_, and demonstrating the emergence of lower probability events than observed in homogeneous tissue. Thirdly, two mechanisms of SCRE mediated conduction block were demonstrated (Figure 4), illustrating mechanisms by which re-entry may be initiated. Finally, the long term interactions with re-entry were investigated (Figure 5-6), demonstrating that sustained re-entrant excitation can load the SR-Ca^2+^ and promote SCRE mediated focal excitation, and revealing a purely functional mechanism of localisation.

### Conditions for the emergence of SCRE mediated focal activity

Analysis of the impact of cellular variability in the dynamics of SCRE on the probability of the emergence of an ectopic beat highlighted the complex considerations surrounding the amplitude and timing of SCRE, and its modulation by *I*_K1_: introducing a proportion of cells more vulnerable to SCRE in combination with reduced *I*_K1_ led to a negative shift the SR-Ca^2+^ relationship as well as the emergence of lower probability events, contrasting to the almost step-function response in homogeneous tissue; the same SCRE variability without reduced *I*_K1_ right-shifted the probability curve, despite the presence of larger amplitude SCRE compared to the homogeneous condition. These data demonstrate the importance of membrane potential and the timing of large scale release: with normal *I*_K1_ there has been insufficient time for the important role of surrounding tissue depolarization to raise the membrane potential and allow the early timed, large amplitude releases to trigger excitation; with reduced *I*_K1_, it is substantially easier for these early timed large amplitude events to overcome electrotonic load. This important role of *I*_K1_, as previously highlighted [10,11,15,19], may be of particular relevance to arrhythmia associated with heart failure remodelling, in which *I*_K1_ is generally reduced.

Furthermore, these lower probability events indicate the presence of a minimal substrate for the emergence of focal activity which may arise with very low probability under conditions without significant single-cell SCRE; certainly, the mechanism by which focal activity emergences permits this possibility: consider the potential rare occurrence associated with low-probability SCRE in which the activity is localised and the large amplitude releases are well timed relative to surrounding tissue to initiate TA. It is possible that these low probability events play an important role in the spontaneous initiation of arrhythmia in predominantly healthy patients. Full characterisation of this minimal substrate is therefore vital for quantitative assessment of the potential importance of these events; such work is currently being carried out.

### Pro-arrhythmic feedback between re-entry and SCRE

The mechanisms by which a suitably timed focal excitation can result in conduction block and the onset of self-perpetuating re-entrant excitation have been extensively studied previously (e.g. [20]); the present study also demonstrates a feedback mechanism by which re-entrant excitation promotes SCRE: Combining simulations of SCRE with those of sustained re-entrant excitation demonstrated that the resulting SR-Ca^2+^ loading can promote the emergence of focal excitation following termination of re-entry, perpetuating the arrhythmic conduction patterns.

The simulations revealed a purely functional mechanism of localised ectopic and re-entrant excitation, without the requirement for a specifically vulnerable region, determined by the additional relaxation time associated with the unexcited core of a scroll wave. This localization combined with rapid focal activity resulted in focal excitation patterns which were highly asymmetric and almost indistinguishable from the re-entry, due to conduction block with the tail of the previous re-entrant or focal excitation.

Furthermore, these asymmetric conduction patterns could result in a complete re-entrant circuit and the re-initiation of sustained re-entry. This mechanism switching may not clearly present in mapping or ECG measurements, but may have significant implications for pharmacological intervention and provide one explanation for variable and limited success. Combination of the approaches presented with biophysically detailed models of healthy and diseased myocardium at the single cell and organ scales will allow detailed investigation of these complex interactions and their potential physiological and pharmacological impact.

### Comparison to other studies

Only a few computational studies have attempted to dissect the multi-scale mechanisms involved in SCRE mediated arrhythmia. Initially, the minimum tissue substrate for the emergence of focal excitations resulting from non-stochastic EADs and DADs was investigated [21], followed by demonstration of independent cellular events emerging as a focal excitation [10–14] and SCRE as a mechanism for both triggered activity and conduction block [15]. Potential interaction with extracellular matrix remodelling was demonstrated [16], as well as the potential non-linear considerations for pharmacological action on both triggers and substrate [13].

Important features observed in these previous studies are in agreement with that of the present study, i.e. in relation to the mechanism of synchronisation overcoming electrotonic load, the steep SR-Ca^2+^ relationship of ectopic activity in homogeneous tissue, the importance of *I*_K1_ in governing the vulnerability to ectopic activity, and the potential role of non-TA inducing DADs to cause conduction abnormalities [10,11,15,19]; independent validation of these features is therefore provided by the present study. The present study also provides novel analysis and mechanistic insight, pertaining to: (i) the analysis of SCRE vulnerability variability on the SR-Ca^2+^-TA relationship; (ii) investigation of the potentially pro-arrhythmic multi-scale coupling between SCRE and re-entry; and (iii) demonstration of a mechanism of localisation of these phenomena which can also lead to focal-re-entrant mechanism switching.

### Limitations

Limitations of the computational framework are discussed in the previous study, Arrhythmia Mechanisms and Spontaneous Calcium Release: I - Multi-scale Modelling Approaches (Under Review). This sections describes only limitations pertaining to the mechanistic analysis presented.

Simulations of SCRE emerging as tissue-scale arrhythmia in general required model parameters and conditions towards the extreme of physiologically observed behaviour (i.e., corresponding to highly diseased myocardium), and therefore do not provide a complete picture of the role of SCRE across the spectrum of patients presenting arrhythmia. Further analysis and new methodological approaches will be required to practically simulate lower probability events and fully assess the role of SCRE in cardiac arrhythmia.

In the wider context, these simulations represent only initial detailed *in silico* analysis of the impact of SCRE at the organ scale: experimental validation, integration with biophysically detailed species and disease specific models, and parameterisation to experimentally measured SCRE statistics, are essential to translate the approaches and mechanistic insight of the present study to arrhythmia in patients.

### Conclusions: From Ca^2+^ sparks to arrhythmia and back

The computational framework applied in this study presents congruent simulations of potentially arrhythmic events from dyad to whole-organ, highlighting the mechanistic pathways and feedbacks which operate from Ca^2+^ spark to arrhythmia (Figure7):

1. Stochastic oscillations of RyRs within a single dyad act to release Ca^2+^ and, promoted by high SR-Ca^2+^ load, initiate a Ca^2+^ spark (“sub-cellular trigger”);
2. The distribution of dyads within the cell provides a dynamical substrate for the propagation of Ca^2+^ sparks as a whole-cell Ca^2+^ wave (“sub-cellular substrate”);
3. Loading the SR-Ca^2+^, such as through rapid pacing under sympathetic activity, can therefore result in the emergence of substantial cellular SCRE and triggered action potentials (“cellular trigger”);
4. Electrotonic coupling plays a dual role in the emergence of focal activity in tissue models, both repressing the effects of SCRE on membrane potential and promoting the spread of small tissue depolarisation (“tissue trigger substrate”);
5. Which allows appropriately timed SCRE to overcome source-sink mismatch and reach the threshold for activation of *I*_Na_ (“tissue trigger”);
6. Non-propagating DADs and rapid focal excitation both present a substrate for the emergence of asymmetric conduction patterns which may degenerate into sustained re-entry (“tissue substrate”);
7. Sustained re-entrant excitation can load the SR-Ca^2+^, promoting SCRE and focal activity (“substrate-trigger feedback”);
8. The interaction of rapid focal excitation with the tail of re-entrant excitation enhances the tissue substrate for conduction block and increases the probability of re-initiation of re-entry and perpetuation of arrhythmia (“trigger-substrate feedback”).

**Figure 7:**
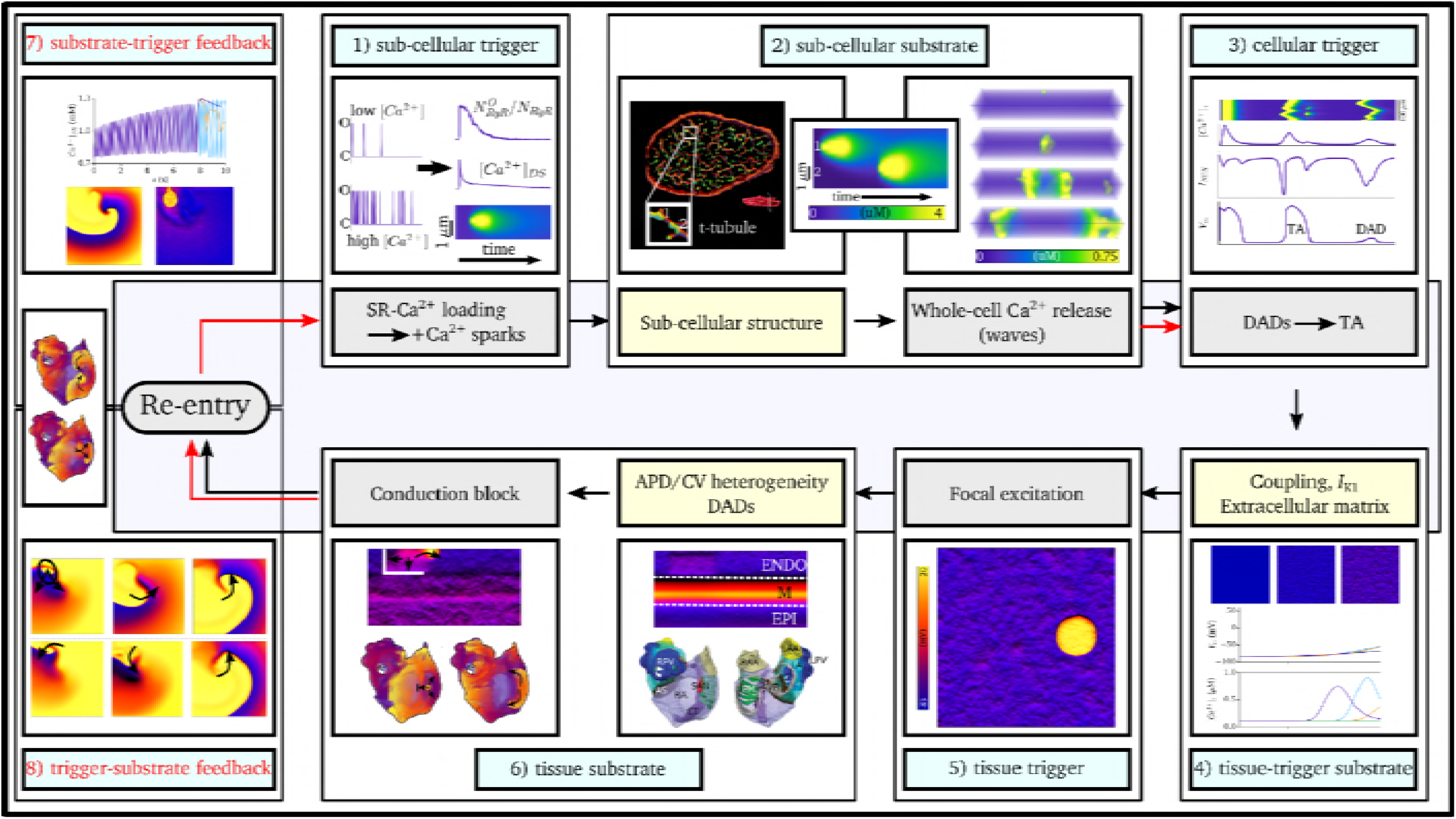
Mechanisms of coupling between SCRE and arrhythmia. Cellular cross section data (part 2) provided by Dr. Izzy Jayasinghe, University of Leeds. Illustration of atrial heterogeneity (part 6) reproduced with permission from [20]. All other illustrations utilise data from this or the related study.

### Final Remarks

By successfully reproducing pro-arrhythmic tissue-scale behaviour mediated by SCRE, results presented in this study and those of other groups [10–13,15,22] support the proposition that stochastic sub-cellular Ca^2+^ dynamics play an important role in arrhythmia initiation and maintenance. However, there still remain many mechanistic and specific questions regarding SCRE and arrhythmia which have yet to be addressed. For example, how can we simulate very rare events which may trigger an arrhythmic episode, and thus explain the role of SCRE in spontaneous arrhythmia initiation? How does the role and importance of SCRE change over the progression of a condition such as heart failure or atrial fibrillation? How does SCRE interact with structural and extracellular matrix factors such as hypertrophy, fibrosis and scar? What role does cellular remodelling and its components (ion channel vs cellular structure) play in the manifestation of SCRE at the organ scale?

## Acknowledgements

I would like to thank Dr Izzy Jayasinghe, University of Leeds, for providing experimental data used in Figure 7, and also to Dr Al Benson for providing comments and feedback on the manuscript.

## Supplementary material

**S1 Text - Model description:** Document containing full model equations and parameters for all of the cell models and SRF presented in this study.

**S1 Code:** C/C++ code containing all implementations presented in the study [Note: will be supplied at revision/acceptance stage following any updates due to review].

**S1 Video - Emergence of focal excitation in tissue:** Intracellular Ca^2+^ concentration (left) and membrane potential (right) illustrating the emergence of a focal excitation. Corresponds to the condition shown in Figure 2.

**S2 Video - DAD mediated conduction block in 2D:** 2D homogeneous sheet, showing the emergence of DADs following regular pacing before an S2 stimulus is applied to the upper edge, resulting in conduction block. Corresponds to Figure 3Aii - upper.

**S3 Video - DAD mediated conduction block in 3D:** 3D homogeneous ventricular wedge, showing the final S1 excitation and an S2 excitation leading to conduction block; both stimuli applied to the ENDO wall. Corresponds to Figure 3Aii - lower.

**S4 Video - Focal mediated conduction block:** Simulation in homogeneous (upper) and heterogeneous (lower) 2D sheets, with focal excitation emerging at a similar time, leading to conduction block only in the heterogeneous case. Corresponds to Figure 3Bii.

**S5 Video - Focal excitation following re-entry pt 1:** Left panels show voltage (upper) and SR-Ca^2+^ (lower) from a randomly selected cell in 2D tissue, right panels show the spatial distribution of voltage (left), intracellular Ca^2+^ (middle) and SR-Ca^2+^ (right). Data show the no SRF case (purple lines; upper 2D panels) and with SRF included (red lines, lower 2D panels), set with the threshold being over but close to the SR-Ca^2+^ peak reached during re-entry. This simulation illustrated a single, not-functionally localised focal excitation.

**S6 Video - Focal excitation following re-entry pt 2:** Left panels show voltage (upper) and SR-Ca^2+^ (lower) from a randomly selected cell in 2D tissue, right panels show the spatial distribution of voltage (left), intracellular Ca^2+^ (middle) and SR-Ca^2+^ (right). Data show the no SRF case (purple lines; upper 2D panels) and with SRF included (red lines, lowest 2D panels), set with the threshold further above the SR-Ca^2+^ than Video S5. This simulation illustrated a single, functionally localised focal excitation.

**S7 Video - Focal excitation following re-entry pt 3:** Left panels show voltage (upper) and SR-Ca^2+^ (lower) from a randomly selected cell in 2D tissue, right panels show the spatial distribution of voltage (left), intracellular Ca^2+^ (middle) and SR-Ca^2+^ (right). Data show the no SRF case (purple lines; upper 2D panels) and with SRF included (red lines, lowest 2D panels), set with the threshold even further above the SR-Ca^2+^ than Video S5 and S6. This simulation illustrated multiple, functionally localised focal excitations.

**S8 Video - Mechanism switching:** AP traces (upper panels) and snapshots of voltage and intracellular and SR calcium (lower panels). A number of re-entrant cycles are shown, followed by rapid focal activity which degenerates back into re-entry.

